# Influence of preprocessing, distortion correction and cardiac triggering on the quality of diffusion MR images of spinal cord

**DOI:** 10.1101/2023.09.26.559530

**Authors:** Kurt G Schilling, Anna J.E. Combes, Karthik Ramadass, Francois Rheault, Grace Sweeney, Logan Prock, Subramaniam Sriram, Julien Cohen-Adad, John C Gore, Bennett A Landman, Seth A Smith, Kristin P. O’Grady

## Abstract

Diffusion MRI of the spinal cord (SC) is susceptible to geometric distortion caused by field inhomogeneities, and prone to misalignment across time series and signal dropout caused by biological motion. Several modifications of image acquisition and image processing techniques have been introduced to overcome these artifacts, but their specific benefits are largely unproven and warrant further investigations. We aim to evaluate two specific aspects of image acquisition and processing that address image quality in diffusion studies of the spinal cord: susceptibility corrections to reduce geometric distortions, and cardiac triggering to minimize motion artifacts. First, we evaluate 4 distortion preprocessing strategies on 7 datasets of the cervical and lumbar SC and find that while distortion correction techniques increase geometric similarity to structural images, they are largely driven by the high-contrast cerebrospinal fluid, and do not consistently improve the geometry within the cord nor improve white-to-gray matter contrast. We recommend at a minimum to perform bulk-motion correction in preprocessing and posit that improvements/adaptations are needed for spinal cord distortion preprocessing algorithms, which are currently optimized and designed for brain imaging. Second, we design experiments to evaluate the impact of removing cardiac triggering. We show that when triggering is foregone, images are qualitatively similar to triggered sequences, do not have increased prevalence of artifacts, and result in similar diffusion tensor indices with similar reproducibility to triggered acquisitions. When triggering is removed, much shorter acquisitions are possible, which are also qualitatively and quantitatively similar to triggered sequences. We suggest that removing cardiac triggering for cervical SC diffusion can be a reasonable option to save time with minimal sacrifice to image quality.

## Introduction

Diffusion magnetic resonance imaging (dMRI) is a medical imaging technique that provides information on the microstructure and anatomic connectivity of the spinal cord (SC). The most commonly used dMRI technique, diffusion tensor imaging (DTI), provides quantitative indices of fractional anisotropy (FA), mean diffusivity (MD), axial diffusivity (AD), and radial diffusivity (RD), which are sensitive to features of tissue microstructure including axonal density, geometry, injury, and degree of myelination [1]. Beyond DTI, several biophysical, or multi-compartment, models of diffusion have been applied to the spinal cord [2-16] providing greater specificity to tissue microstructure and greater insight into biology or pathology.

Despite these promises, SC dMRI on human subjects is challenging due to SC size and location [17]. The SC is a thin structure, requiring relatively high spatial resolution for adequate image contrast. Also, the cord is surrounded by various moving tissues which can introduce artifacts from local susceptibility variations and motion. Several modifications of image acquisition and image processing techniques have been introduced to overcome these artifacts while maintaining adequate contrast and resolution. This study investigates two strategies to address dMRI artifacts: susceptibility correction to mitigate geometric distortions with image processing, and cardiac triggering to mitigate motion artifacts during image acquisition.

### Distortions in spinal cord EPI data

Diffusion MRI is typically collected using echo planar imaging (EPI) [18]. EPI suffers from susceptibility artifacts which manifest as distortions in the phase-encoding (PE) direction of the image. This is especially prominent in the cord which experiences large B0 field variations due to proximity to the lungs, strong CSF flow, and alternating bone-disc-bone tissue structure.

Preprocessing methods to correct distortions include mapping the B0 field which can be used to estimate and correct voxel-wise displacements in the image [19]. A more common strategy is to acquire pairs of images with reverse PE directions, often referred to as a “blip-up blip-down” acquisitions, yielding pairs of images with equal but opposite distortions. These data can be used to estimate a distortion field which brings the images to the mid-way point between them (i.e., the undistorted image) [20]. However, these preprocessing techniques have largely focused on, and been optimized for, brain imaging, and there is much work needed to understand optimal pipelines for analyzing SC dMRI. Few studies have investigated this in the SC. Evaluating a variety of reverse PE algorithms in the cervical SC, Snoussi et al. [21, 22] find that distortion correction algorithms provide small, but significant, improvements in both alignment of diffusion tensors along the cord and geometric similarity to structural images.

Similarly, Dauleac et al., [23] show that this improved alignment of diffusion tensors after distortion correction improves tractography quality. Inspired by these works, we aim to evaluate the effects of distortion correction in the SC. We extend these analyses by investigating different acquisition strategies (varying phase encoding directions, number of diffusion-encoding directions) and different preprocessing strategies for distortion correction across different segments of the cord (cervical and lumbar), in both healthy controls (HC) and subjects with multiple sclerosis (MS).

### Cardiac Triggering

The SC also experiences substantial physiological motion. This includes cardiac motion, respiratory motion, and cerebrospinal fluid pulsation, which cause translation in the superior-inferior and anterior-posterior directions and a possibly nonlinear compression or stretching of the cord [24, 25]. Motion artifacts may cause signal or slice-dropout artifacts, misalignment of diffusion images, and subsequent biases in diffusion quantification [24]. To potentially reduce motion-related artifacts, it has become standard practice to use cardiac triggering to acquire data at a phase of the cardiac cycle where motion is minimal [17]. However, triggering dramatically increases scan time, which can be prohibitive when high angular resolution or multi-shell experiments are necessary for multi-compartment diffusion modeling. Towards this end, our second aim is to evaluate the impact on image quality of removing cardiac triggering. Specifically, we hypothesize that advances in preprocessing and fitting procedures may overcome motion-related artifacts, removing the need for cardiac triggering, and gaining significant scan time savings without an increase in artifacts. To do this, we designed experiments to directly test differences with and without triggering, and assess image quality, image artifacts, and changes in quantitative parameters and reproducibility. Finally, we examined whether time saved by removing triggering can be used to acquire more diffusion weighted images without loss of image quality.

## Methods

### Distortion Correction – data acquisition

Data to evaluate distortion correction included seven cohorts (**Table 1**): 4 HC cervical cohorts, 1 MS cervical cohort, 1 HC lumbar cohorts, and 1 MS lumbar cohort. Local Institutional Review Board approval and written informed consent were obtained prior to imaging. Imaging data for all participants was acquired using 3.0T whole-body MR scanners (3T dStreamine Ingenia and 3T Elition X, Philips Achieva, Best, Netherlands). A two-channel body coil was used for excitation and a 16-channel SENSE neurovascular coil was used for reception. DTI sequences were acquired with single-shot EPI using a reduced field-of-view (FOV) outer volume suppression technique [26] in the axial plane at an effective b-value of 750 s/mm^2^, including a b=0 s/mm^2^ volume and 15 diffusion weighted images in uniformly distributed directions. The protocol included a cardiac-gated spin-echo acquisition with SENSE factor = 1.5, flip angle = 90°, TR = 5 beats (∼5000 msec), TE = 70 msec, resolution = 1.1 × 1.1 mm^2^, slice thickness = 5 mm, number of slices = 14, centered on C3/C4 vertebral level, FOV = 64 × 48 mm, number of acquisitions averaged (in k-space) =3. Additionally, a high-resolution (0.6 × 0.6 × 5 mm^3^) multi-slice, multi-echo gradient echo (mFFE) [27] image was acquired (TR/minTE/ΔTE = 700/7.2/8.8 msec, α = 28°, number of slices = 14, 3 echoes) over the same volume in order to act as an undistorted structural image for registration and subsequent comparison with diffusion data.

**Table 1.** Data used to investigate susceptibility distortions in SC diffusion MRI includes different SC segments, phase encode directions, diffusion weighted directions, and cohorts.

| Dataset | Segment | Phase Encode | Directions | b-value | Sample size | Cohort |
| --- | --- | --- | --- | --- | --- | --- |
| Cervical | Cervical | RL | 15 | 750 | 55 | HC |
| Cervical MS | Cervical | RL | 15 | 750 | 54 | MS |
| Cervical #2 | Cervical | RL | 15 | 750 | 16 | HC |
| Cervical AP | Cervical | AP | 32 | 750 | 10 | HC |
| Cervical RL | Cervical | RL | 32 | 750 | 10 | HC |
| Lumbar | Lumbar | RL | 15 | 750 | 28 | MS |
| Lumbar MS | Lumbar | RL | 15 | 750 | 4 | MS |

While the above-described sequence was considered the base sequence, all cohorts had small differences. Cervical #1 (N=55 total datasets, 42 subjects, 13 with two visits; mean age ± standard deviation = 31.6 ± 7.1 years) and Cervical MS (N=54 total datasets, 40 subjects, 14 with two visits; mean age ± standard deviation = 34.7 ± 6.5 years) followed this sequence exactly for HC and MS subjects, respectively, for projects involving the study of SC diffusion in MS (NIH R01NS117816). Cervical #2 included HC for a project investigating stenosis of the cord (NIH R01NS104149) and used the same acquisition but was centered lower at the C4/C5 vertebral level and consisted of an older cohort (mean age ± standard deviation = 53 ± 9.4 years). Cervical AP and Cervical RL images were acquired to investigate effects specifically of phase encode direction (and consequently distortion direction) (NIH K01EB032898), with phase encoding in the AP and RL directions, respectively. These cohorts also included over double the amount of diffusion directions (32 versus 15). Finally Lumbar and Lumbar MS datasets were acquired as part of a project to optimize protocols in the lumbar cord (NIH K01EB030039). For Lumbar datasets a dual channel body coil (transmission) and a 12-channel in-couch SC array (receive) were used. Data were centered at the SC lumbar enlargement (corresponding to vertebral levels ∼T11-L1), with resolution, slices, b-values, gradient directions, TR (5 beats), and number of images averaged all remaining the same as in cervical SC imaging.

### Distortion correction – preprocessing

Preprocessing was performed following 4 pipelines, all of which started with a diffusion denoising performed on the raw, acquired dataset using the Marcenko-Pastur PCA algorithm [28] implemented in MRTrix3 (v3.0.4) [29] (*dwidenoise*). The first included no motion correction nor distortion correction (referred to as *RAW*) and served as a baseline for which to compare improvements in image quality. The second pipeline (*MOCO*) included motion correction only, using the Spinal Cord Toolbox (v5.8) [30] algorithm (*sct_dmri_moco*) inspired by [31], which uses a slice-wise registration based framework to align diffusion weighted volumes with regularization along the rostral-caudal axis (polynomial of order 2), and a group-wise registration across time series (3 groups used here). The third pipeline (*TOPUP*) used the FSL (v6.0.6.1) software package [32] using the *topup* and *eddy* algorithms [20] which used the reverse PE pair to correct motion, susceptibility distortion, and eddy currents. We zero-padded the first and last slices (2 slices padded to each) to ensure that the algorithm included these during the processing pipeline. The fourth pipeline was the Hyper-elastic Susceptibility artefact Correction method [33] (*HySCO*) (v2.0), which was implemented as part of the ACID toolbox and similarly uses a physical distortion model to best match the forward and reverse phase encode images.

Following preprocessing, the diffusion tensor [34] was fit voxel-wise to the outputs of each preprocessing pipeline using standard least squares fitting (Spinal Cord Toolbox, *sct_dmri_compute_dti*). Next, for every preprocessing pipeline, the b=0 image was registered to the structural mFFE using a slice-wise center of mass alignment of the cord segmentation, followed by rigid registration using Mutual Information between the images as the registration minimization metric (*sct_register_multimodal*). Finally, nonlinear registration between the mFFE and the PAM50 spinal cord template [35] (*sct_register_to_template*) was performed, which enabled the propagation of tissue type labels (white matter and gray matter) and vertebral labels to each preprocessed image (code repository at https://github.com/schillkg/SpinalCord-Distortion-Triggering). WM and GM of the lumbar cohorts were manually labelled.

### Distortion correction – evaluation

First, a qualitative evaluation was performed by visualizing the b=0 image, the mean diffusion weighted image (*meanDWI*), and the FA map rigidly registered to the anatomical mFFE. This enabled visual comparisons of size, shape, and asymmetry of the diffusion data as well as potential artifacts in the data or introduced by the processing pipelines.

We then performed two analyses to quantitatively compare diffusion indices (b=0, meanDWI, FA, AD) to the mFFE image. First, we calculated the normalized mutual information (NMI) between each contrast and the mFFE (which acts as a ‘ground truth’ geometry without geometric distortions) - using the full field of view (i.e., a 40mm window around the center of the cord that includes all tissue) to calculate NMI (using the Pattern Recognition and Machine Learning Toolbox v1.0.0.0 in MATLAB R2021_b). Next, to assess whether distortion corrections truly improve the intra-cord alignment or are simply driven by CSF and surrounding contrasts, we calculated NMI between each contrast and the mFFE but limited to the cord alone using the SC segmentation derived from the mFFE. For both the full field-of-view and cord-only field of view, comparisons between the 4 distortion-correction preprocessing choices were performed using a paired-sample t-test to assess statistically significant differences in NMI between paired observations (i.e., different preprocessing algorithms on the same dataset) correcting for multiple comparisons (4 algorithms x 7 datasets x 4 quality metrics).

To further assess distortion correction capability within the cord, we calculate the contrast to noise ratio (CNR) between white matter (WM) and gray matter (GM) tissues. CNR was calculated as the contrast between tissues (WM minus GM) divided by the noise calculated as the standard deviation of metric intensity within the WM. We chose contrasts of FA and AD, as these are expected to show strong WM/GM contrast. Because the WM and GM masks are derived from the ground truth structural image, any misalignment or asymmetries due to distortions will likely result in a reduced CNR.

Finally, we assessed the variability in the derived DTI indices as a function of distortion correction algorithm to evaluate whether the choice of correction method impacted the derived indices. To do this, we calculated the averaged values of FA and AD within WM and GM respectively, and again tested for differences between pipelines using the paired-sample t-tests.

### Cardiac Triggering – data acquisition

Our experimental protocol includes 6 cervical SC acquisitions (**Table 2**), each repeated twice within a scan session. First, we used the diffusion acquisition from the Spinal Cord Generic protocol (SCG) [17, 36] – a recently introduced protocol intended to provide a harmonized acquisition of high-quality MRI of the cervical cord, regardless of site or vendor. Second, we used the protocol described in the methods above (referred to as VUIIS). Both protocols use cardiac triggering to acquire dMRI data during the quiescent phase of cardiac-related motion. While both SCG and VUIIS dMRI acquisitions are closely matched for TE/TR, differences in these acquisitions include number of diffusion directions, number of signals averaged, different reduced field of view techniques, and phase encoding directions. For each protocol (SCG and VUIIS), we also acquired a matched protocol with no triggering (SCG-nt; VUIIS-nt), with all other acquisition parameters matched (where the “estimated” scan duration is in theory equivalent). Next, we acquired a time-matched VUIIS protocol without triggering and without averaging, enabling the acquisition of 64 diffusion directions in an equivalent time (VUIIS-nt-time), from which we can select a subset of 15 directions to match the original protocol (VUIIS-nt-dir) which has an equivalent scan time of 1 minute.

**Table 2.**
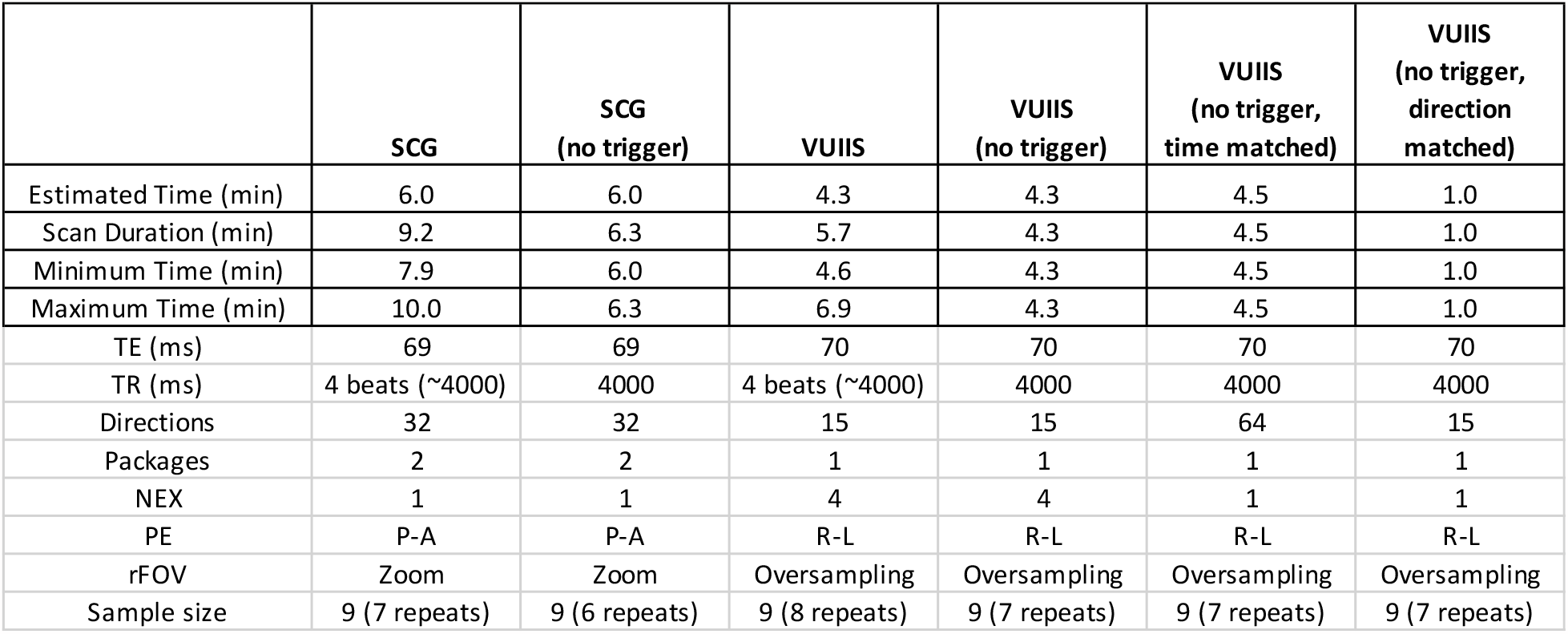
Scan protocols for cardiac triggering experiments. Triggering experiment included the Spinal Cord Generic (SCG) protocol with and without triggering, VUIIS protocol with and without triggering, a VUIIS protocol without triggering or signal averaging, and a 1-minute subset of this protocol. NEX is number of excitations, or averages. Overall, cardiac triggering increases scan time.

|  | SCG | SCG<br>(no trigger) | VUIIS | VUIIS<br>(no trigger) | VUIIS<br>(no trigger,<br>time matched) | VUIIS<br>(no trigger,<br>direction<br>matched) |
| --- | --- | --- | --- | --- | --- | --- |
| Estimated Time (min) | 6.0 | 6.0 | 4.3 | 4.3 | 4.5 | 1.0 |
| Scan Duration (min) | 9.2 | 6.3 | 5.7 | 4.3 | 4.5 | 1.0 |
| Minimum Time (min) | 7.9 | 6.0 | 4.6 | 4.3 | 4.5 | 1.0 |
| Maximum Time (min) | 10.0 | 6.3 | 6.9 | 4.3 | 4.5 | 1.0 |
| TE (ms) | 69 | 69 | 70 | 70 | 70 | 70 |
| TR (ms) | 4 beats (~4000) | 4000 | 4 beats (~4000) | 4000 | 4000 | 4000 |
| Directions | 32 | 32 | 15 | 15 | 64 | 15 |
| Packages | 2 | 2 | 1 | 1 | 1 | 1 |
| NEX | 1 | 1 | 4 | 4 | 1 | 1 |
| PE | P-A | P-A | R-L | R-L | R-L | R-L |
| rFOV | Zoom | Zoom | Oversampling | Oversampling | Oversampling | Oversampling |
| Sample size | 9 (7 repeats) | 9 (6 repeats) | 9 (8 repeats) | 9 (7 repeats) | 9 (7 repeats) | 9 (7 repeats) |

### Cardiac Triggering – preprocessing

Based on the results of the distortion correction experiments, we chose to use only motion correction in the cardiac triggering experiments, although the preprocessing steps remained largely the same. For each dataset, we used MRTrix [29] for denoising, followed by Spinal Cord Toolbox [30] for motion correction, tissue segmentation, and metric extraction, as described previously. One difference here is that we used the RESTORE algorithm [37] for outlier-robust tensor fitting.

### Cardiac Triggering – evaluation

Qualitative evaluation across protocols was first performed by visualization of individual DWIs and FA maps for each protocol. Second, artifact prevalence with/without triggering was assessed by quantifying several QA measures from the SCILPY toolbox (https://github.com/scilus/scilpy) including the % of slice signal dropouts, mean residuals from a tensor fit, physically implausible signals, and pulsation/misalignment derivations. Third, changes or bias in derived indices due to triggering were quantified by calculating the FA and MD in white and gray matter at the C3-C4 level with and without triggering. Fourth, reproducibility was quantified by calculating a % Error from scan-rescan measures of the same metrics, calculated as a measure of difference divided by mean value. Significant differences between triggered and non-triggered protocols (for QA measures, changes in measures, and reproducibility measures), as well as between direction- and time-matched protocols were determined using paired t-tests with multiple comparison correction.

## Results

### Distortion Correction

Qualitative results are shown in **Figure 1**, with a single example image shown for each dataset (additional examples are shown in Supplementary Documentation). Here, the b=0, mean DWI, and FA maps are shown for RAW, MOCO, TOPUP, and HYSCO algorithms. Clear distortions are present in the raw data, corresponding as expected in the phase encoding direction. These distortions exist within the MOCO data, with little observable differences in derived measures. Both TOPUP and HYSCO visibly correct distortions, most apparent in the b=0 images where the CSF surrounding the SC is both left-right symmetric and qualitatively matches that from the anatomical image. However, little differences are observed in either the mean DWI or FA image. Importantly, a smudging/smearing artifact is often observed when performing distortion correction (both TOPUP and HYSCO), most apparent in the mean DWI.

**Figure 1.**
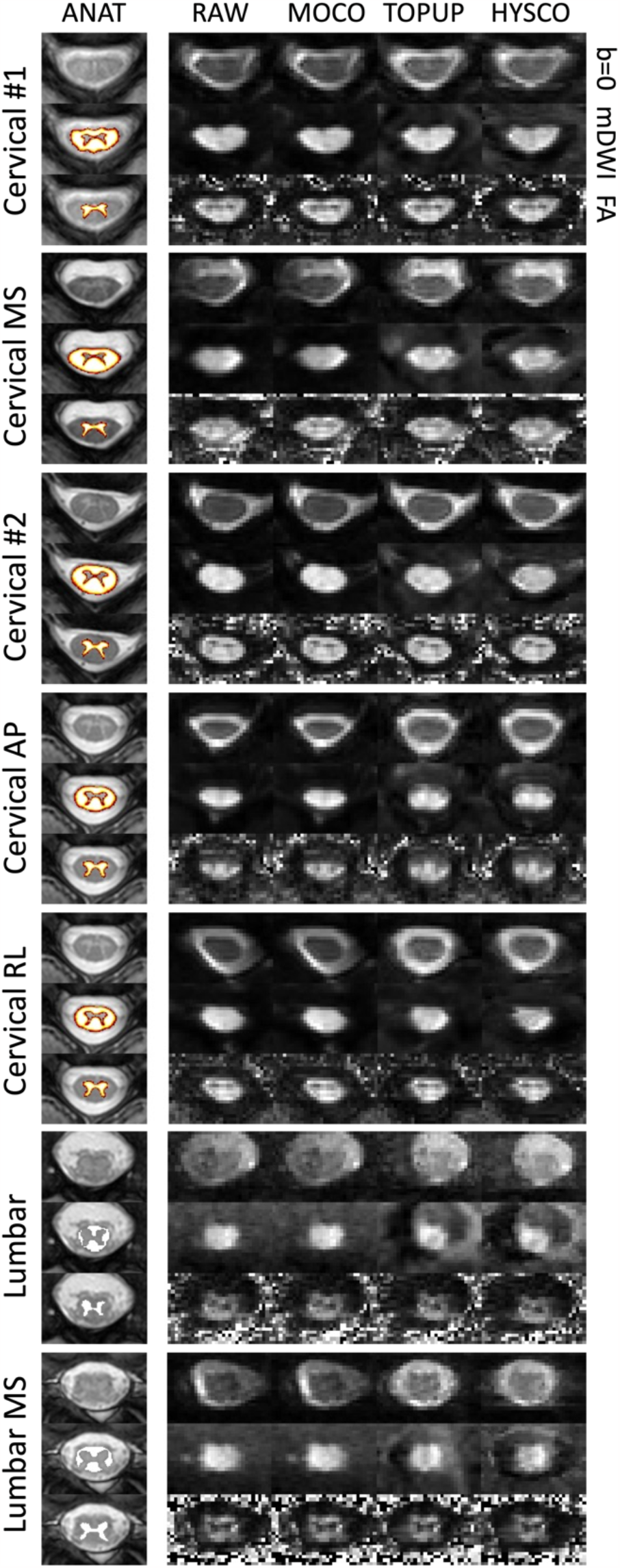
Distortion correction algorithms qualitatively correct susceptibility distortions in the spinal cord, however, artifacts may be introduced within the diffusion-weighted images and subsequent DTI maps. The b=0, mDWI, and FA map of an example subject from each of the 7 cohorts is shown after no correction (RAW), motion correction only (MOCO), or correction using the TOPUP or HYSCO algorithms. Middle axial slice is selected for visualization. Note that WM and GM labels are probabilistic for cervical cohorts but binary for lumbar cohorts (due to manual labelling).

**Figure 2**. shows quantitative results of the NMI between the anatomical image and diffusion contrast (b=0, meanDWI, FA, AD), for all cohorts and pipelines. For several contrasts, particularly meanDWI and AD, the NMI is increased with motion correction, and further increased with both TOPUP and HYSCO algorithms – with the two distortion correction algorithms performing comparably. While it is challenging to compare across datasets due to different anatomies, structural sizes, and image contrasts, there is generally lower NMI in lumbar versus cervical cord, lower NMI in the aging cohort (C2), and lower NMI in RL phase encoded versus AP.

**Figure 2.**
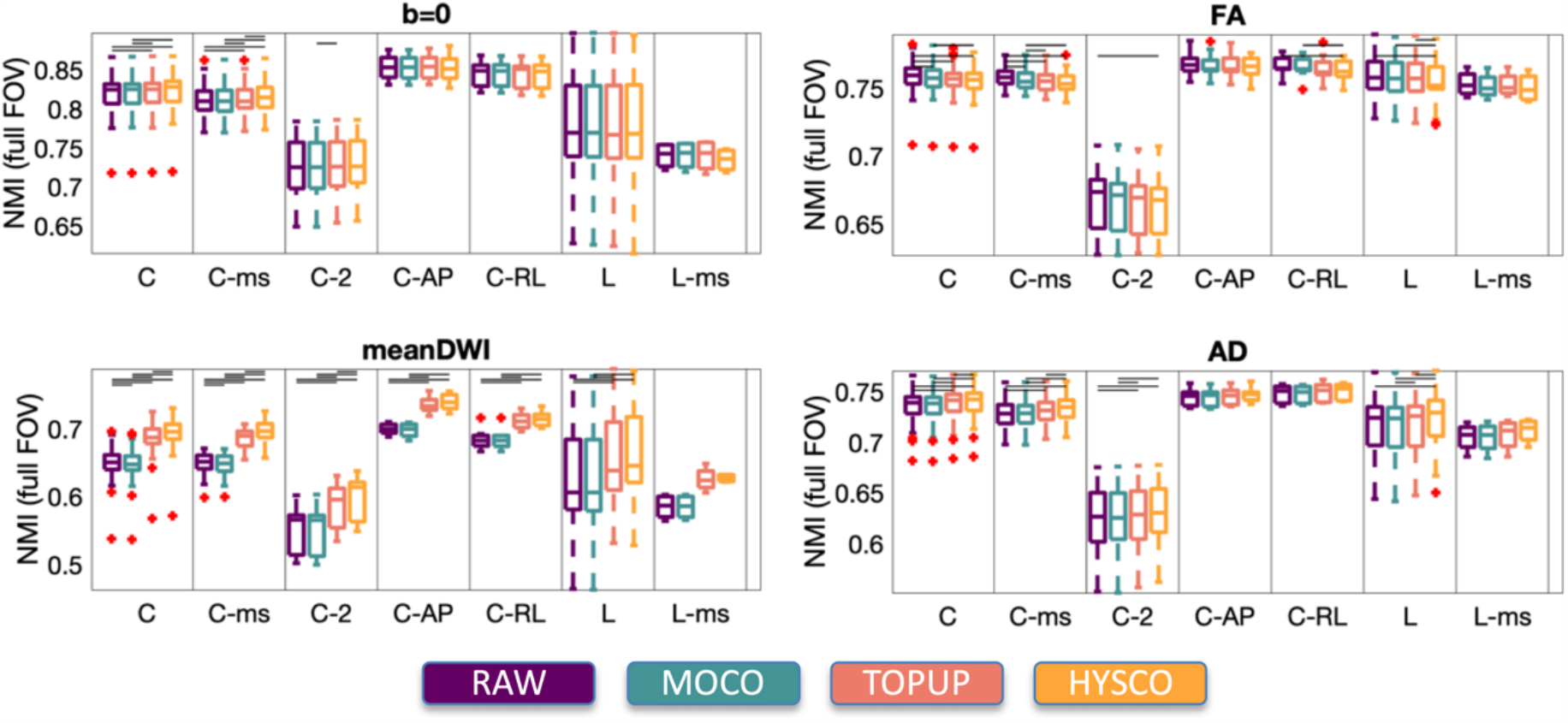
Motion correction and distortion correction algorithms improve NMI between anatomical images and diffusion contrasts. NMI over the full image field of view is calculated between diffusion contrasts (b=0, meanDWI, FA, AD) and the mFFE contrast and is shown for all cohorts. Horizontal bars indicate statistical significance after multiple comparison correction.

**Figure 3**. shows quantitative results of the NMI between the anatomical image and diffusion contrast but calculated within the cord only. In contrast to the full field-of view results (**Figure 2**), several metrics do not improve significantly (FA, AD), or even get worse with distortion correction (b=0), although the meanDWI shows improvements in most datasets. However, improvements in meanDWI similarity to mFFE are significantly attenuated compared to those assessing the full field of view (with CSF).

**Figure 3.**
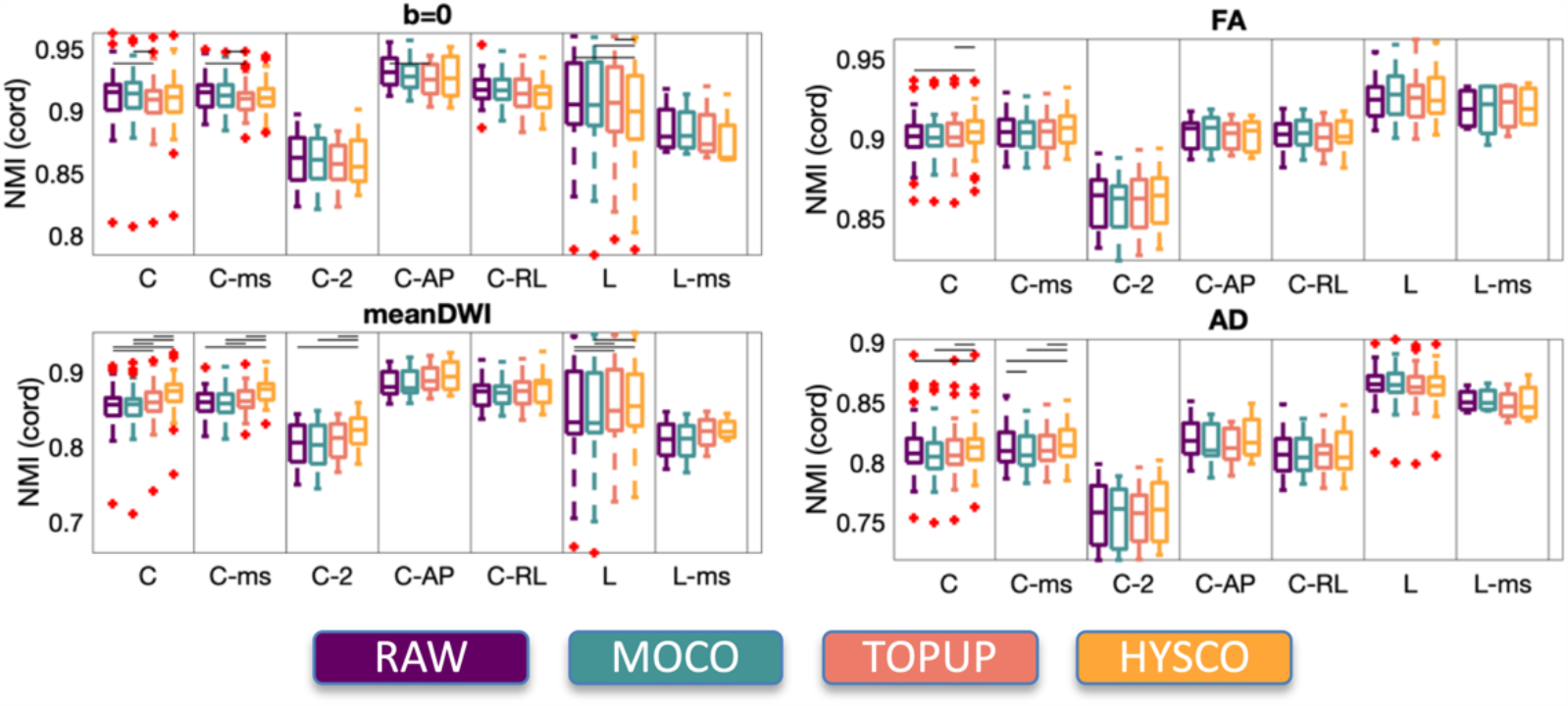
Improvement in alignment to the structural image is attenuated when assessing cord structure only. NMI calculated within the SC between diffusion contrasts (b0, meanDWI, FA, AD) and the mFFE contrast and is shown for all cohorts. Horizontal bars indicate statistical significance after multiple comparison correction.

**Figure 4**. shows CNR between white and gray matter for FA and AD, two DTI measures which are expected to reflect contrast between these tissue types. There is little statistically significant improvement in AD CNR due to motion/distortion correction in any cohort. However, while not always reaching statistical significance, motion correction does increase FA CNR in both cervical and lumbar segments in controls and MS patients. Similarly, TOPUP does increase FA CNR, particularly compared to no correction at all.

**Figure 4.**
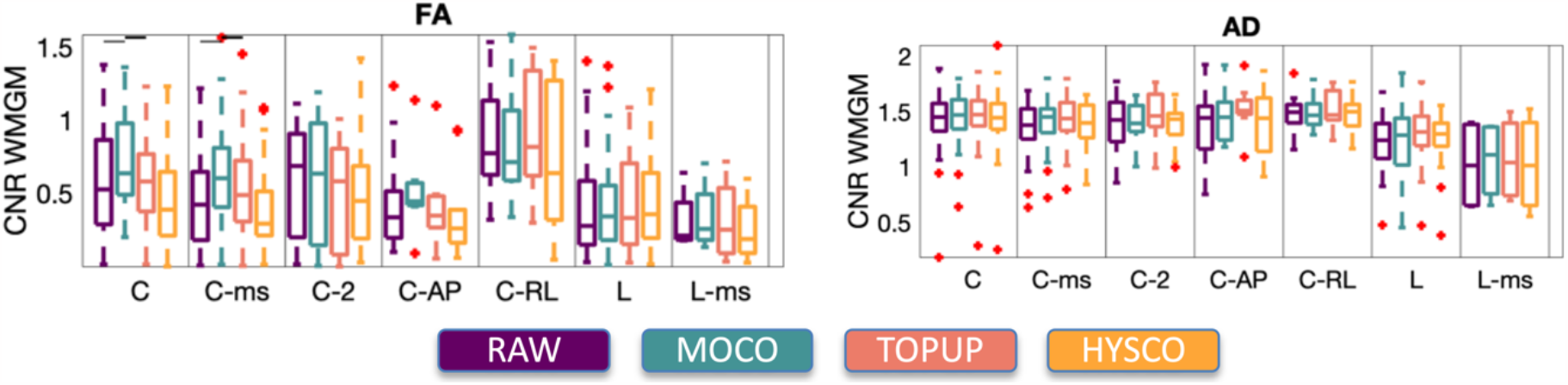
Motion correction improves CNR between white and gray matter tissues. Because WM/GM CNR is related to image quality and artifact attenuation, motion correction only (MOCO) generally outperforms other algorithms (or no processing at all) – however, the improvement is small and not always statistically significant.

**Figure 5**. shows that motion correction and distortion correction change the DTI-derived indices in WM and GM, in expected ways. In the WM, these image preprocessing steps generally increase WM FA and AD - both of which are expected to be higher in white than in gray matter – although, again, the change is often not statistically significant. In many cases, distortion correction, both TOPUP and HYSCO, decrease WM FA relative to motion correction only.

**Figure 5.**
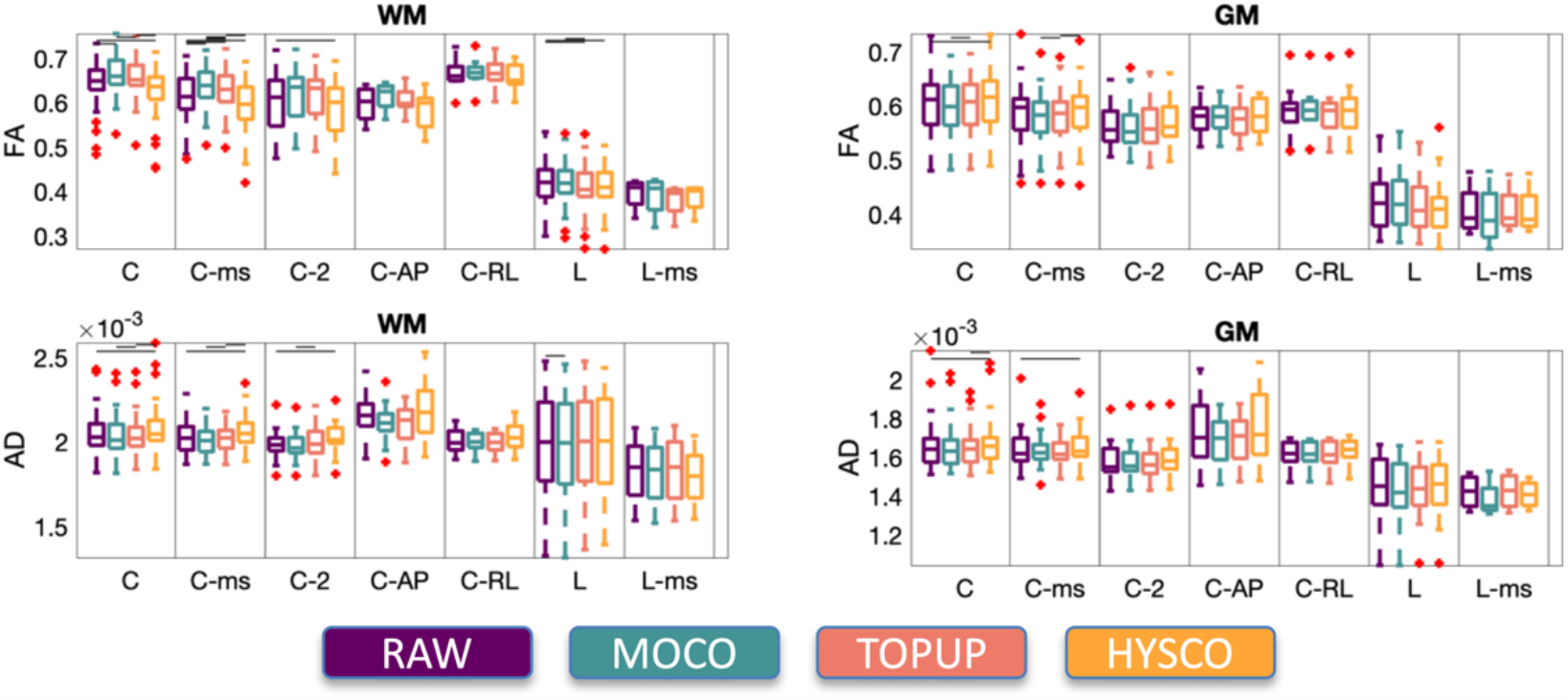
DTI measures do change after motion and distortion correction in the spinal cord. FA and AD of the WM and GM are shown for each cohort, with statistically significant changes between preprocessing pipelines indicated by a horizontal bar.

### Cardiac Triggering

**Table 2** shows the estimated and true scan durations of each protocol, both triggered and non-triggered. While the estimated scan durations are matched, the true duration increases on averaged 46% and 32% with triggering for SCG and VUIIS protocols, respectively.

**Figure 6**. shows the middle slices for a single diffusion image and the derived FA map for an example subject for each acquisition protocol. Visually, very little differences are observed between triggered/non-triggered, and similar results are observed with the 1-minute reduced protocol (**Figure 1**, column 6).

**Figure 6.**
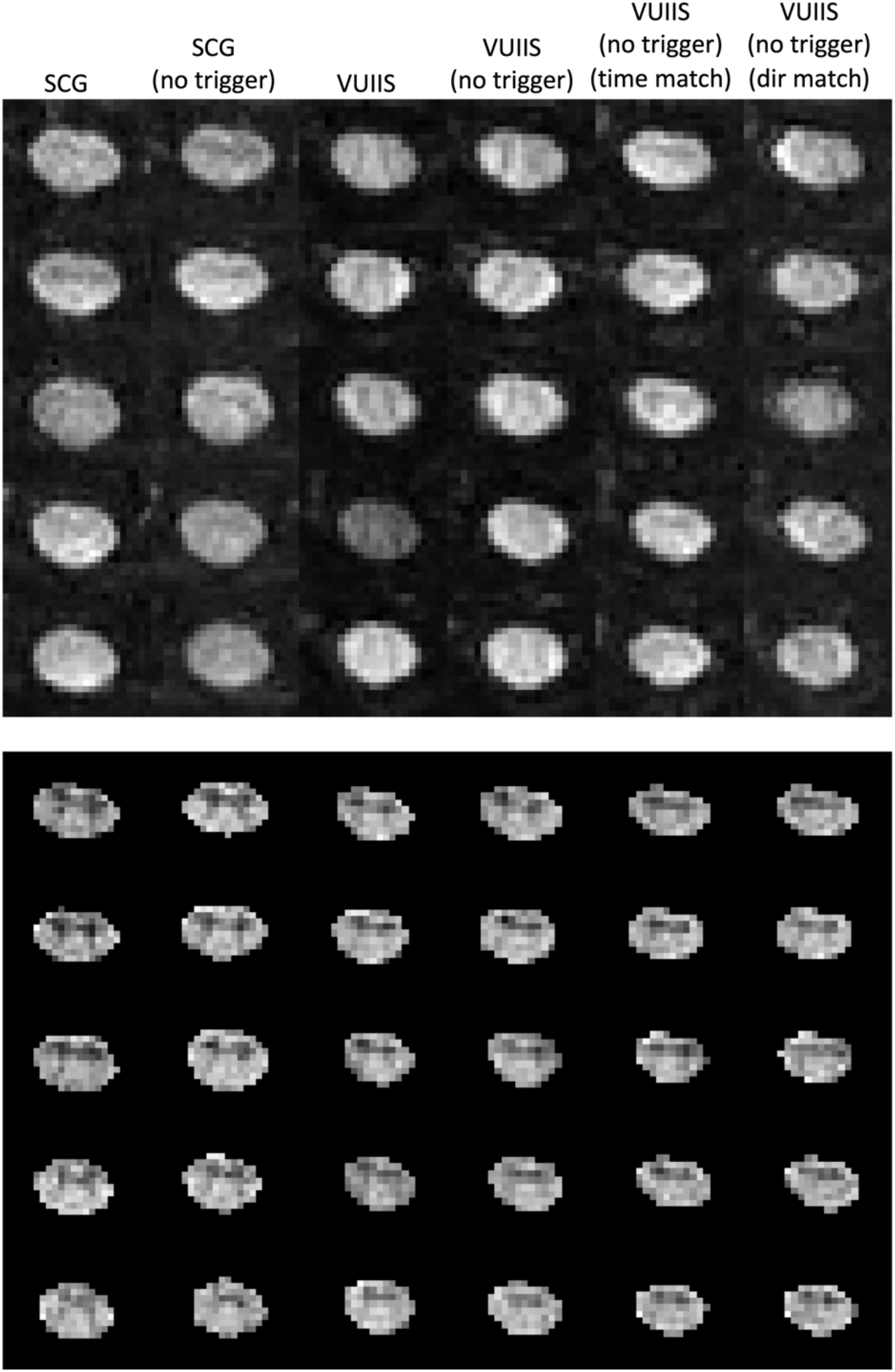
Qualitative comparison of acquisitions (DWI top; FA bottom) show very little differences between triggered and non-triggered acquisitions. Similar results are observed for the time-matched and direction-matched acquisitions which allow more directions or shorter time, respectively.

**Figure 7**. plots quality control measures for each acquisition protocol. For both SCG and VUIIS protocols, there is no significant increase in any artifacts, residuals, or misalignment when removing triggering. However, there are increased residuals with the time-matched 64 directions sequence, likely due to the decreased SNR from decreased averaging.

**Figure 7.**
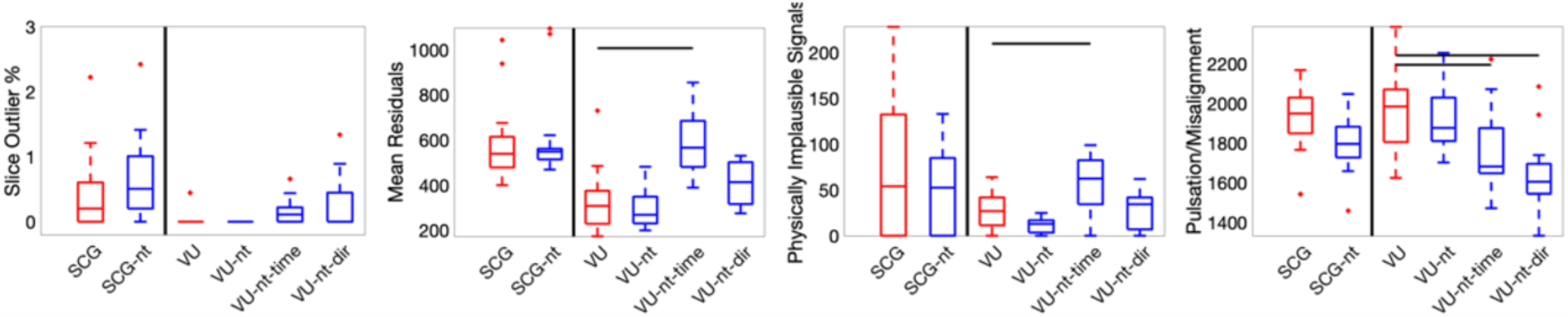
Quality control metrics show no statistically significant differences between trigger/non-triggered. However, differences exist between triggered and the direction or time-matched protocols.

**Figure 8**. shows the scan rescan measures for all protocols with and without triggering.

**Figure 8.**
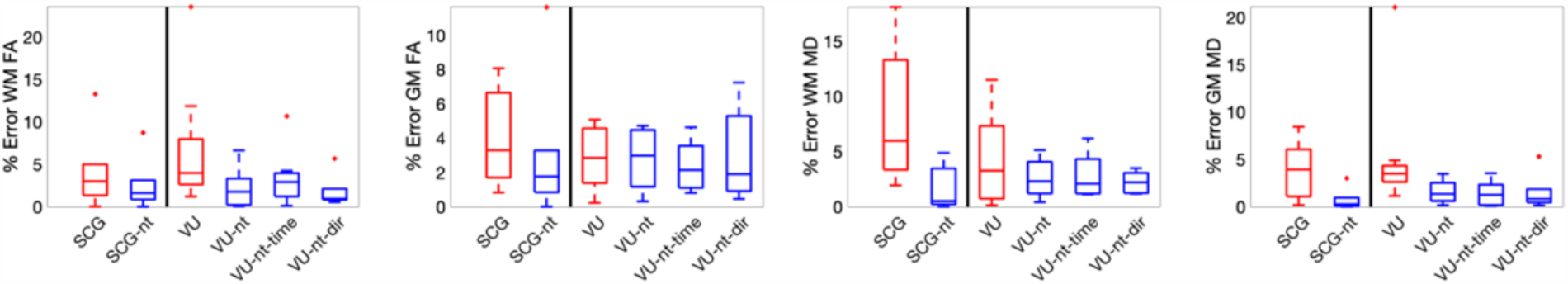
Reproducibility is quantified as a measure of % error between scan and rescan for FA and MD in WM and in GM. No significant differences in reproducibility exist between triggered and non-triggered protocols.

The scan rescan reproducibility of derived indices does not significantly increase due to the removal of triggering. In fact, the reproducibility decreases in many cases, although the decrease is not statistically significant.

Finally, to show that results generalize to a patient population, the triggering experiments for a clinical MS protocol were repeated on a small sample (N=5) of MS patients. In agreement with previous findings, triggering increased scan time by ∼33% compared to the non-triggered acquisition (**Supplementary Documentation**), DWIs and FA maps are qualitatively similar between triggered and non-triggered acquisitions (**Figure 9**), artifacts are not significantly increased (**Supplementary Documentation**), and DTI-derived measures are not significantly different with and without triggering (**Supplementary Documentation**).

**Figure 9.**
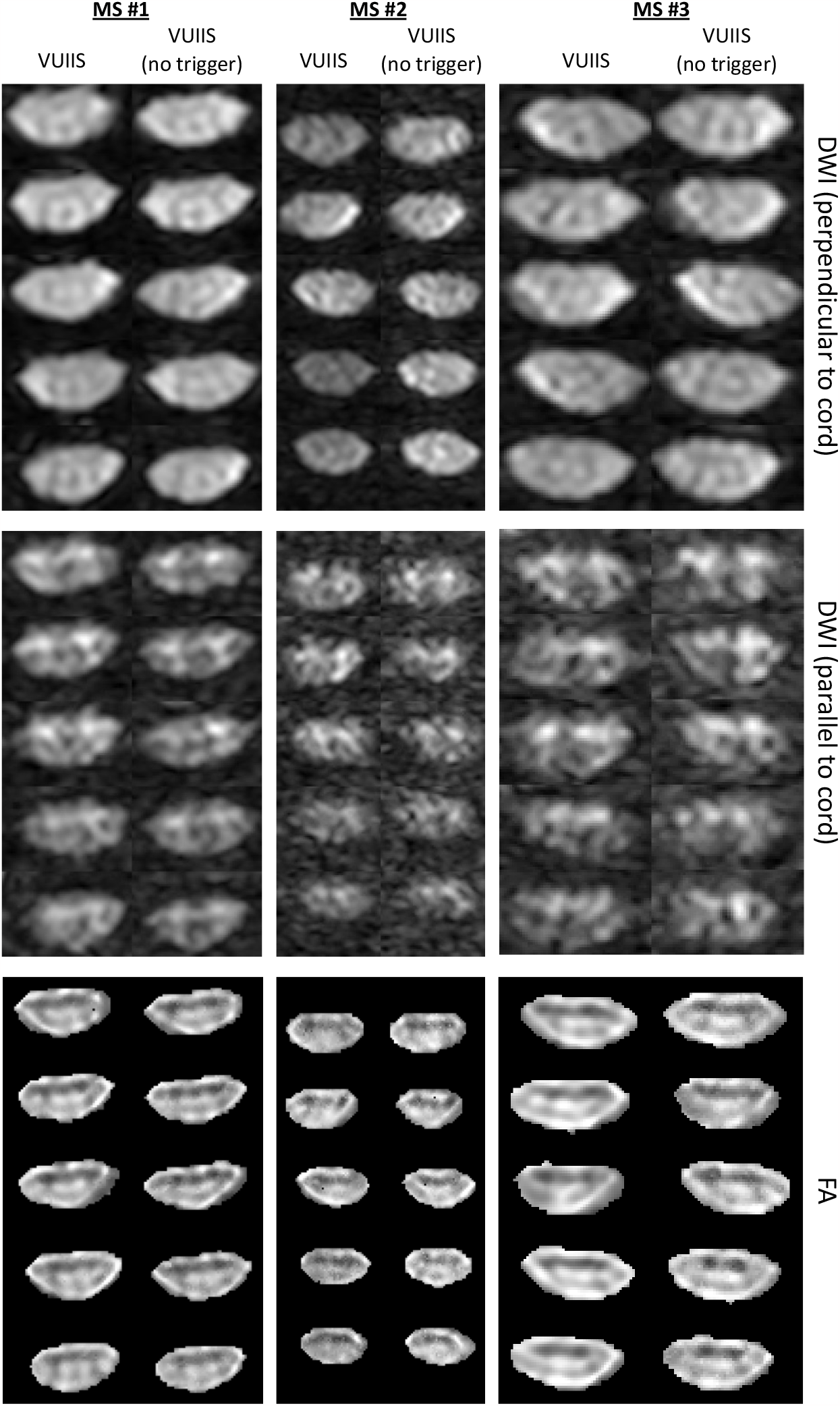
Little qualitative differences between cardiac triggered and non-triggered in an MS cohort. Consecutive slices of the diffusion image most perpendicular to the cord, most parallel to the cord, and an FA map are shown for three MS patients.

## Discussion

### Distortion correction

Diffusion MRI of the SC suffers from susceptibility artifacts resulting in geometric distortions of the images. Accurately correcting distortions facilitates better alignment with structural images and more reproducible quantitative MRI indices, together resulting in more precise quantitative description of the SC tissue types or detailed regions of interest. While several techniques have been proposed to address distortions [20, 38-41], they have largely been developed and optimized for brain diffusion MRI, and their efficacy in the cord has not been fully validated.

Using 7 datasets of the cervical and lumbar SC, we compared and evaluated 4 possible preprocessing strategies that use reverse phase encoded data to correct distortions. We find that distortion correction techniques increase geometric similarity to structural images, but this improved similarity is largely driven by the high-contrast CSF, and the geometry within the cord is not always consistently corrected to match that of the structural image. In fact, often we see unnatural blurring or smudging artifacts, particularly in the diffusion weighted images or in the DTI-derived quantitative maps. Based on these results, our recommendation is that, at the very least, bulk motion-correction must take place in the preprocessing pipeline.

These results generalize across acquisitions, cohorts, and disease state – with similarities between cervical and lumbar observations. Results are also in alignment with previous studies investigating distortion correction on isotropic imaging voxels over a larger field of view [21, 22]. Snoussi et al. [21] presented a comparison of 4 different distortion correction methods, finding no significant difference in DTI-based metrics pre/post correction, but did find improved local alignment of diffusion tensors in parallel with the superior/inferior direction of the cord. In supplementary experiments we find improved alignment of the cord (**Supplementary Documentation**), but with statistically significant improvement in the largest cervical dataset only. Further supplementary analysis showed that most significant distortion occurred at the C3/C4 vertebral level in the cervical cord, with patterns less clear in the lumbar cord, but greater distortion/misalignment near T10/T11 vertebral levels. An additional supplementary finding was that distortion magnitude is not related to curvature of the cord (**Supplementary Documentation**).

Adaptations and improvements to current distortion preprocessing algorithms are needed, as most are currently designed and optimized for brain imaging. As discussed in the forum [42] for the Spinal Cord Toolbox [30], one reason for the substandard performance of reverse phase encode algorithms could be due to the reduced field of view techniques used in imaging this structure, in combination with flow artifacts, that cause a mismatch in the information content of images and hinders these algorithms (which expect exactly equal and opposite distortions). Beyond this, assumptions (and default parameters) of movement, expected field variations [43], and anisotropic structure (particularly in the current study) may reduce performance of these algorithms. In parallel with preprocessing improvements, changes and innovations in acquisitions may be what are primarily relied on to reduced distortions – including these reduced field of view techniques [44], multi-shot imaging [45-49], or improved shimming and scan set up [17].

There are several limitations to these distortion experiments. Measures of mutual information, CNR, diffusion metric change do not fully characterize geometric fidelity after distortion correction, but instead paint a picture that provides insight into effects of different correction methods. Many measures may be sensitive to other artifacts (ghosting, fat shift, movement) beyond distortion alone. The registration is also extremely sensitive to the algorithm and parameter choices. We have chosen to perform a simple center of mass alignment plus image-based rigid-registration to not alter the geometry of the diffusion measures when comparing to the undistorted structural image. Even though RAW and MOCO approaches result in more geometric discrepancies with the mFFE, these discrepancies could be addressed by the registration step through nonlinear registration, which is not investigated in this study. Finally, diffusion and mFFE contrasts are expected to represent different features of the tissue, and alternative similarity metrics could have been chosen to evaluate similarity to the structural image.

### Triggering

Cardiac triggering is commonly used in dMRI protocols of the cervical, thoracic, and lumbar cord to reduce cardiac-related motion artifacts. However, this can dramatically increase scan time which can limit studies requiring more diffusion directions or higher b-value data, for multi-compartment diffusion modeling, for example. Here, we use two different (but typical) diffusion protocols in the cervical cord to investigate the effects of removing cardiac triggering from the acquisition protocol. We hypothesized that removing triggering will not only shorten scan time but also result in qualitatively and quantitatively similar as cardiac triggered acquisitions.

First, we confirmed that triggering greatly increases scan time in our investigated protocols. For both protocols, the scan time using cardiac triggering was ∼30-50% longer than a simple scan time calculation of Repetition Time x Number of Excitations (i.e., the time of the non-triggered acquisition). The potential increase in scan time is dependent on TR (number of beats), number of slices, and resolution/bandwidth, and is not expected to be the same for all sequences. Second, minimal qualitative differences were observed between triggered and non-triggered acquisitions. Raw images, mean DWIs, and DTI-derived indices are qualitatively similar with and without triggering. Third, quality control metrics showed no significant differences either – with no increase prevalence of slice dropouts, residuals, nor misalignment artifacts.

Fourth, scan rescan reproducibility also did not decrease when triggering was removed. And finally, the results generalized to a patient population. Overall, we suggest that cardiac triggering can be removed from the acquisition process without sacrificing image quality, which enables shorter scan times and/or more diffusion weighted images.

Most SC dMRI studies use cardiac triggering, motivated in part by several early studies showing motion driven largely by respiration, followed by cardiac-related effects [50-52].

Translation was observed to be greatest in the superior-inferior and anterior-posterior directions and on the order of ∼1mm, varying along the length of the cord and varying throughout the cardiac cycle [25]. Validation in physical phantoms confirmed that signal dropout can be caused by this motion [24], although the dropout is likely due to the non-rigid compression and stretching of the cord rather than pure translation in any direction. However, few studies have directly measured and compared image quality with and without triggering.

We hypothesized that removing cardiac triggering may not be overly detrimental to the diffusion-weighted images not because the SC doesn’t move, but because artifacts caused by these motions may be overall minor, and somewhat reconcilable with advances in image processing (e.g., denoising) and model fitting (e.g., robust outlier detection and down-weighting). Our results confirm this hypothesis, as the images themselves are not appreciably different (i.e., artifacts not overly prevalent) and the derived indices are also not different (i.e., if outliers exist, they do not change final DTI fit). It is important to emphasize that these results may not generalize at all times, particularly pathology where increased motion has been observed in cervical myelopathies [53, 54] and stenosis [55], and similar quality assurance should be performed prior to removing triggering for a given study.

Beyond removing triggering, we show that replacing signal averaging with acquisition of more, unique, diffusion directions does not hinder subsequent analysis. Acquiring more diffusion images has the added benefit of enabling high angular resolution imaging methods or advanced modeling beyond the simple tensor model. Further, a simple, 15 direction, single-average, and non-cardiac-triggered acquisition (approximately 1 minute acquisition) results in DTI indices similar to not only the 4-average non-cardiac-triggered acquisition but also the 4-average cardiac triggered acquisition. These results are in line with early DTI studies of the brain suggesting ∼20+ unique directions are necessary for robust estimation of anisotropy and diffusivity [56, 57]. In summary, removing triggering can be used for either scan time savings or acquisitions of more unique diffusion directions, diffusion times, or diffusion weightings.

## Conclusion

Using several study cohorts, in different SC segments, and with different acquisition conditions, we investigated two specific aspects of SC diffusion MRI: distortion correction and cardiac triggering. We recommend at a minimum to perform bulk-motion correction in preprocessing and find that improvements are needed in distortion correction pipelines for consistent, artifact-free analysis in this organ. Second, we show that removing cardiac triggering allows significant scan time reduction without sacrificing image quality and suggest that this scan time savings can be used for acquisition of more diffusion weightings or diffusion directions.

## Acknowledgements

NIH K01EB030039 (KO), K01EB032898 (KS), 5R01NS109114 (SS), 5R01NS117816 (SS) and 5R01NS104149 (SS), R01EB017230 (BL), R01NS104149 (JG).

